# Dynamics of miRNA-driven feed-forward loop depends upon miRNA action mechanisms

**DOI:** 10.1101/016162

**Authors:** Maria Duk, Maria Samsonova, Alexander Samsonov

## Abstract

**Background:** We perform the theoretical analysis of a gene network sub-system, composed of a feed-forward loop, in which the upstream transcription factor regulates the target gene via two parallel pathways: directly, and via interaction with miRNA.

**Results:** As the molecular mechanisms of miRNA action are not clear so far, we elaborate three mathematical models, in which miRNA either represses translation of its target or promotes target mRNA degradation or is not re-used, but degrades along with target mRNA. We examine the feed-forward loop dynamics quantitatively at the whole time interval of cell cycle. We rigorously proof the uniqueness of solutions to the models and obtain the exact solutions in one of them analytically.

**Conclusions:** We have shown that different mechanisms of miRNA action lead to a variety of types of dynamical behavior of feed-forward loops. In particular, we find that the ability of feed-forward loop to dampen fluctuations introduced by transcription factor is the model and parameter dependent feature. We also discuss how our results could help a biologist to infer the mechanism of miRNA action

## Background

An important role in regulation of gene expression in higher eukaryotes, plants and animals belongs to miRNAs, the endogeneous small non-coding RNAs that bind to partially complementary sequences in target mRNAs. The miRNAs are involved in regulation of development, differentiation, apoptosis, cell proliferation [1, 2, 3, 4], as well as in the progression of numerous human diseases, such as chronic lymphocytic leukemia, fragile X syndrome, and various tumor types [5, 6, 7, 8, 4]. Each miRNA molecule may target hundreds of mRNAs, and, vice versa, some targets are combinatorially affected by multiple miRNAs [9, 10, 11]. Comparative phylogenetic studies uncovered the conserved miRNA-binding sequences in more than one third of all genes, that lead to a suggestion that the miRNA regulation may be relevant to a large portion of cellular processes [12, 13].

However, it should be noted that there is no common notion about mechanisms of miRNA action. There are experimental evidences that miRNAs regulate gene expression through translational repression, mRNA deadenylation and decay, however the contribution and timing of these effects remain unclear [14, 15]. Although some studies show translational repression without mRNA decay [16], others point to decay as a primary effect [17, 18]. It was also demonstrated in several papers that the destabilization of target mRNA is accompanied by degradation of a miRNA molecule [19, 18].

It is well known that transcription networks contain several biochemical wiring patterns called network motifs. One of the most significant recurring motifs is the feed-forward loop (FFL) [20, 21, 22], in which the upstream transcription factor (TF) regulates the target gene via two parallel pathways: directly, and by interaction with a second molecule, which also regulates the target gene. An assigned value for a pathway is defined as positive, if the total number of negative interactions in the pathway is even, and negative otherwise. The FFL is named as “coherent” if the sign of *direct* regulation path coincides with the overall sign of the *indirect* regulation path, and “incoherent” otherwise.

Recent computational analysis demonstrated that FFL, containing TF and miRNAs, are overrepresented in gene regulatory networks, assuming that they confer useful regulatory opportunities [23]. There are two possible structure configurations in each coherent and incoherent FFL, containing miRNA, as shown in Figure 1. We will specify these configurations as the type 1 or 2 coherent FFLs, and the type 1 or 2 incoherent FFLs, respectively.

**Figure 1.**
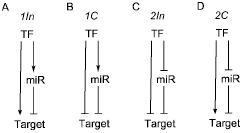
The incoherent and coherent feed-forward loops. Arrows mean activation, the turned over T-bars indicate repression. TF -transcription factor, miR -miRNA, Target - target protein. **A**: 1In - type 1 incoherent FFL, TF activates both target mRNA and miRNA synthesis. **B**: 1C - type 1 coherent FFL, TF represses traget mRNA and activates miRNA synthesis. **C**: 2In - type 2 incoherent FFL, TF represses both target mRNA and miRNA synthesis. **D**: 2C - type 2 coherent FFL, TF activates target mRNA and represses miRNA synthesis.

The role of miRNA in gene regulatory networks becomes currently a subject of wide speculations. In general, it is expected that FFLs with miRNA can buffer the consequences of noise action in gene expression in order to confer a robustness to environmental perturbation and genetic variation [12, 24]. The anti-correlative expression of miRNAs and their target mRNAs was documented in many cases [25, 12], where it points out that while transcription primarily controls target gene expression, the miRNAs lend further reinforcement to gene regulation by attenuation any unwanted transcripts. The second class of genetic buffering by miRNA emerges in cases, where both the miRNA and the target are coexpressed at intermediate levels [26, 27, 28]. It was proposed that intermediate miRNA quantities and low target avidity (the result of a single “seed” binding site) intentionally provide the target protein synthesis, but place a burden on this process, which, most likely, provides an approach to buffer stochastic fluctuations in the mRNA level [12]. However, it should be noted that up to date there is the only direct observation that miRNA buffers gene expression against perturbation [29].

A simple mathematical modeling was used recently to explore the capability of FFLs with miRNA to buffer fluctuations in gene expression. Osella et al. introduced the model [30], describing a target gene regulation in the type 1 incoherent FFL, however, solved analytically the coupled *algebraic* equations only, obtained by trivial reduction of the coupled 1st order ordinary differential equations (O.D.E.) to a steady state. For stochastic statement of the problem in [30] O.D.E. were solved numerically. It was shown that with respect to the simple gene activation by TF, the introduction of the miRNA-mediated repressing pathway can significantly dampen fluctuations in the target gene output for essentially all the choices of input parameters and initial conditions. Moreover, this noise buffering function was expected to be direct consequence of the peculiar topology of the FFL.

There is a critical imperfection in any mathematical model based on the steady state data analysis. Firstly, recent measurements [31, 32] of protein abundance and turnover by parallel metabolic pulse labeling for more than 5,000 genes in the NIH 3T3 mouse fibroblasts showed that the half live times of many proteins in these cells are longer than the cell cycle duration of 27.5 hours. For proteins, which half life time is comparable or longer than the cell cycle, the genuine steady state is not reached before the cell division. As a consequence, quantities per cell are not only defined by synthesis and degradation, but also by an *initial number* of protein molecules at the beginning of cell cycle. Secondly, an absence of time derivatives (as it is in the steady state approximation) is provided by simple replacement of the initial coupled kinetic non-linear equations with corresponding algebraic equations, avoiding any information about the temporal behavior of cell components. A variety of the cell process models usually demonstrate versatile dynamics at early stages even in case of coincidence of stationary values. For these reasons it is much more faithful and informative, however difficult, to study coupled ODEs at every time moment, aiming to decipher mechanisms underlying its dynamics.

Another challenge in mathematical modeling of biological systems consists in *the non-uniqueness* of solutions to model equations governing a biological problem. This non-uniqueness is partly caused by imperfections of current experimental methods, which are often unable to provide the measurements of all the parameters required to describe the system dynamics. As a result the parameter values have usually to be found by a numerical fitting of the solutions to data. This task, called “an inverse problem”, is, in a way, tricky, because there are usually several parameter sets in good agreement with data. It means that within a given accuracy there are several models, which are, in a sense, equivalent in the data description. This non-uniqueness provided by numerical simulations represents a crucial problem for contemporary mathematical modeling in biology, and usually an appropriate solution may be found by invoking additional information from complicated experiments. On the other side, an intrinsic non-uniqueness may have transparent biological reasons, as living systems are robust and can tolerate large parameter variation, as long as the core network topology is retained [33, 34]. Therefore, mathematical treatment and refinements are still required to better understand how the component interactions result in the system complex behavior.

Exact mathematical solutions to biological problems are free of many imperfections mentioned, however, most of problems are quite complex and cannot be treated analytically without significant simplifications. A few successful quantitative solutions to problems of mathematical biology are known, that do not require the parameter fitting. These solutions are, in fact, the exact solutions to corresponding differential equations. Also, it is worth to mention the importance of analytical solutions, which are valid for every parameter and coefficients set, and, for this reason, provide a genuine ‘check point’ for any numerical simulation obtained by means of different methods.

Here we consider a gene network sub-system composed of a FFL mediated by TF and miRNA. We perform the analysis of three mathematical models, which describe the dynamics of gene expression in FFL under an assumption of different mechanisms of miRNA action. In contrast to recent considerations [30] we examine the FFL dynamics quantitatively at the whole time interval of cell cycle. We rigorously proof the uniqueness of solutions to these models and obtain the exact solutions in one of them analytically. We show how different mechanisms of miRNA action lead to distinctive dynamical behavior of FFLs. In particular, we find that the ability of FFLs to dampen fluctuations introduced by TF is the model and parameter dependent feature. We also discuss how our results could help a biologist to infer the mechanism of miRNA action.

## Results and Discussion

### Mathematical models of FFLs

The biological system under study will be described with 5 variables, representing the number *w* of mRNA molecules transcribed from the TF gene, the number *q* of TF molecules, the number *s* of miRNA molecules, the number *r* of mRNA molecules transcribed from the target gene, and the number *p* of target protein molecules.

We shall use the models proposed in [30] and consider 4 conventional configurations of FFLs with miRNA (see Figure 1. For each gene in FFL we consider transcription, translation, degradation and interactions between genes in the regulatory network.

As molecular mechanisms of miRNA action are not clear so far we consider three different models:

- the model, in which miRNA represses translation of its target – *Stop model*,
- the model in which miRNA promotes target mRNA degradation –*Target degradation model* and
- the model in which miRNA is not re-used but degrades along with target RNA –*Dual degradation model*.

where the ‘working titles’ for the models introduced above will be used for brevity.

Thus we write, the five coupled differential equations proposed in (1) for a feedforward loop (FFL).

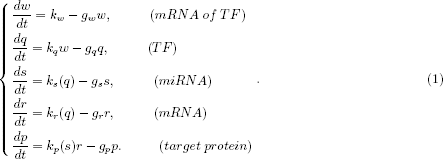

Here *k*_*w*_ and *k*_*q*_ are rates of TF mRNA and TF synthesis, *k*_*s*_(*q*) and *k*_*r*_(*q*) are rates of transcription of the regulated gene, *k*_*p*_(*s*) is the rate of target protein synthesis; *g*_*w*_, *g*_*q*_,*g*_*s*_,*g*_*s*_ and *g*_*p*_ represent the degradation rates of the corresponding species.

Before to specify the types of production functions in (1-4) let us remind that various dynamic processes in a complex system can be described as a progression from an initial quantity that accelerates (or decelerates in case of repression) and approaches a plateau over time. When a detailed description is lacking, a sigmoid function is used, that is based on an idea by A.V. Hill (1910) to describe the sigmoidal *O*_2_ binding curve of haemoglobin. The general form of the Hill function is written as:

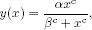

where both *α, β* are arbitrary given parameters of the process. The rate *c* represents either production (*c* ≥ 2) of a quantity *x* or repression of it (*c* ≤ –2) in time. Consequently, the Hill function will be represented by either uprising or falling down graph, and its slope depends on the value of *c*. The easiest way to get a non-trivial regulation type is to prescribe the rate values *c* = 2 and *c* = –2 for activation and repression, respectively, however, even in this case the mathematical problem becomes the non-linear one.

In our problem the production functions *k*_*s*_*(q)* and *k*_*r*_*(q)* are assumed to be the classical Hill functions in the form *k(q) = (k_max_q^c^)/(h^c^ + q^c^)*. The parameters *h*_*s*_ and *h*_*r*_ specify the amount of TF, at which the transcription rate of the miRNA or target gene reaches one half of its maximal value (*k*_*s*_ or *k*_*r*_), and a number *c* is the Hill coefficient, representing the steepness of the regulation curve, see expressions (2) below.

Therefore for the type 1 coherent FFL (see Figure 1) the Hill functions will have the following form:

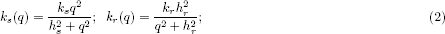

while for type 1 incoherent FFL the production function for target mRNA will be different:

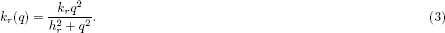

The difference between two types of each of coherent or incoherent loops is in form of the production functions for both target mRNA and miRNA (see Figure 1).

We shall consider three different mechanisms of the miRNA action, following a formalism introduced in [30]. To model the effect of direct translational repression of target mRNA we consider the translation rate of the target *k*_*p*_*(s)* to be a repressive Hill function of the number of miRNA molecules: *k_p_(s) = (k_p_s^c^)/(h^c^ + s^c^)*. The parameter *h* specifies the quantity of miRNAs, at which the translation rate reaches one half of its maximal value *k*_*p*_, and *c* = –2 is again the Hill coefficient.

To model miRNA action in the destabilization of target mRNA we add to the basal rate of mRNA degradation *g*_*r*_ (in absence of miRNAs), a term, which represents an increasing Hill function of a copy number of miRNAs, where *g*_*max*_ is the maximal value of the degradation rate in case of high miRNA concentration, *h*_*deg*_ is the dissociation constant of miRNA-mRNA interaction, and *c* = 2 is the Hill coefficient.

Following [30], we shall discuss also a tentative destabilization of target mRNA, accompanied by the miRNA degradation. The miRNA forms a complex with its target, which degrades with it, instead of being re-used. This complex degradation constant will be denoted as *k*_*rs*_, and the resulting non-linear (due to the multiplicative term *rs*) equations will have a form:

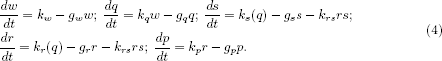

Details of mathematical analysis of the coupled ODE are given in Additional file 1. For the *Stop* model we obtained the exact solutions to the non-linear O.D.E., which coefficients explicitly depend on initial values. Main advantage of exact solutions to corresponding non-linear differential equations is that they do not require any parameter fitting and provide also a reliable basis for verification of numerical results. For the *Target* and *Dual* degradation models the numerical solutions to the problem are obtained. For each model considered we proved also the uniqueness of solutions, i.e., the one-to-one correspondence between given parameters and solutions.

### Comparative analysis of FFL temporal behavior under different models

The mechanisms underlying miRNA-mediated repression are not clear so far, and for this reason we consider three models of the miRNA action described in section.

Four different topologies of FFLs mediated by TF and miRNA are possible in theory, as in Figure 1. We shall analyse the dynamical behavior of all these network topologies in frame of the models described above, that leads, in total, to consideration of 12 different variants of regulation in FFLs.

Further we shall use for brevity the following abbreviations for FFL identification: 1*C* will mean the type 1 coherent loop, 1*In* - type 1 incoherent loop, 2*C* - type 2 coherent loop and 2*In* - type 2 incoherent loop, resp.

We assume that the initial number of molecules in a loop is equal to one half of that obtained just before cell division, which often (however, not necessarily) corresponds to the steady state level. The results for the Stop model are based on exact and explicit solutions obtained to the non-linear O.D.E. 1 (see Additional file 1), and, therefore are free of any data fitting procedure and numerical approximations. For this reason exact solutions may provide also a reliable base and “check points” for numerical solutions in those (Target and Dual degradation) models, where an exact solution is hardly obtainable.

We begin with brief description of the temporal variation of the molecule quantity of each player and in each of three models. To simplify comparison we shall use one and the same parameter set (see section) for all FFL and models.

#### Stop model

In the Stop model the behavior of target mRNA and protein is different in all FFLs because miRNA stops translation of the former and does not promote its degradation. Both 1*C* and 2*In* loops form the identical bell-shaped target mRNA profiles due to repression by TF, in both 1*In* and 2*C* loop the target mRNA profiles are also identical and increase in time to a steady state value (Figure 2). In both 1*In* and 1*C* loop, in which TF activates miRNA gene, the target protein shows pulse-like behavior due to repression mediated by miRNA (Figure 2A,B,E,F). However at steady state in the 1*C* loop the number of target protein molecules is much lower than in the 1*In* loop.

**Figure 2.**
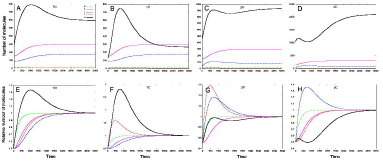
The solutions to the Stop model in various FFLs. 1In - type 1 incoherent FFL, 1C-type 1 coherent FFL, 2In - type 2 incoherent FFL, 2C - type 2 coherent FFL; *w*, *q*, *s*, *r* and *p* denote graphs of solutions for TF mRNA, TF, miRNA, target mRNA and target protein correspondingly. **A - D**: temporal dynamics of absolute number of each molecule species is presented. **E - H**: molecules numbers for each species are normalized on steady state values to better visualize the behavior of RNA species.

In 2*C* loop the number of target protein molecules firstly slightly rises, than falls down due to miRNA action), but with continuous repression of the miRNA synthesis, the target protein begins to rise up to the steady state level (Figure 2H,D).

In 2*In* loop the target protein profile also exhibits the *wave-like* behavior (Figure 2C,G). Firstly, it rises to almost stationary level, after that it slightly decreases due to repression of mRNA synthesis by TF and repression of mRNA translation by miRNA, and eventually grows again to the stationary level due to decrease of the miRNA molecules number.

#### Target degradation model

Contrary to the Stop model in the Target degradation model the temporal dynamics of both target mRNA and protein is similar. In this model the target protein production is a linear function of the target mRNA molecules number, and miRNA promotes the degradation of target mRNA. In 1*C* loop two mechanisms are active, namely, mRNA degradation under miRNA action, and the repression of target mRNA synthesis. In 2*In* loop TF represses miRNA production. This explains the difference in dynamics of target RNA and protein in these loops (Figure 3B,F,C,G). In both loops the dynamics shows *pulse-like* behavior, however in the 1*C* loop the numbers of target mRNA and protein molecules are smaller (about 1300 vs. 1500 molecules) at steady state, and the steady state level is reached later (about 4000 sec vs. 3000 sec) in comparison with 2*In* loop.

**Figure 3.**
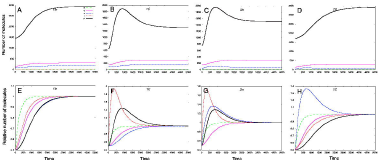
The solutions to the Target degradation model in various FFLs. 1In - type 1 incoherent FFL, 1C- type 1 coherent FFL, 2In - type 2 incoherent FFL, 2C - type 2 coherent FFL; *w, q, s, r* and *p* denote graphs of solutions for TF mRNA, TF, miRNA, target mRNA and target protein correspondingly. **A - D**: the temporal dynamics of absolute number of each molecule species is presented, **E - H**: molecules numbers for each species are normalized on steady state values to better visualize the behavior of RNA species.

The dynamics of target mRNA and protein in both the 1*In* and the 2*C* loops has a form of *increasing* function, tending to a constant value (Figure 3A,D,E,H). The loops behavior differs in time, when the steady state levels of the target mRNA and protein are reached, as well as in numbers of target mRNA and protein molecules at steady state. In 1*In* loop these numbers are smaller (about 3000 molecules vs. 3500 molecules) than in 2*C* loop, that can be explained by promotion of the target mRNA degradation by miRNA in the 1*In* loop.

#### Dual degradation model

In this model both miRNA and target mRNA degrade due to the duplex formation, and the miRNA molecules are not re-used. As in the Target degradation model both target mRNA and protein show similar dynamical behavior. The dynamics of target mRNA and target protein both in 1*C* and 2*In* loops shows *pulse-like* behavior (shown in Figure A1 of Additional file 2), similar to that observed in these loops in the Target degradation model (as shown in Figure 3B,C,F,G). In this model in 2*In* loop the steady state level is approached earlier (3000 sec vs 4000 sec), than in 1*C* loop. The target protein molecules number at steady state in 2*In* loop is also higher than in 1*C* loop (1800 molecules vs 1400 molecules). Moreover, all these numbers in the Dual degradation model are higher than in Target degradation model (Figure A1 of Additional file 2).

In both 1*In* and 2*C* loops the dynamics of target mRNA and protein takes a form of *increasing* function, tending to constant value (Figure A1 of Additional file 2). As in target degradation model in 1*In* loop the steady state is reached earlier (2500 sec vs. 4000 sec) and the level of target protein at steady state is lower (3000 molecules vs. 4000 molecules) than in 2*C* loop. Again, in the Dual degradation model these numbers are bigger than in the Target degradation model.

### Dynamical behavior of FFLs

In this section we shall describe the behavior of each type of FFL in the framework of each model under consideration. We shall demonstrate that FFLs mediated by miRNA and TF may have many possible outcomes, depending on the nature of the relationships between the loop elements.

The study of dynamical behavior of FFLs is based on the variation of initial conditions, and model coefficients with subsequent analysis, how these changes affect the target protein molecules number. To simplify comparison in each numerical experiment we use one and the same parameter set, described in the) section, except of the coefficient value, which effect on the loop behavior is analyzed.

The efficiency in control of the target protein synthesis in all three models depends upon the quantity of TFs (which, in turn, is a function of *k*_*w*_ and *k*_*q*_), the number of miRNA copies (i.e., the function of *k*_*s*_ and *h*_*s*_ defining the affinity of TF to the promoter of miRNA gene), as well as the strength of the miRNA action. In the Stop model the strength of repression of mRNA translation by miRNA is defined as 1*/h*_*p*_. In the Target degradation model the degradation of the target mRNA is described by a term, which represents an increasing Hill function of a copy number of miRNAs, while 1*/h*_*g*_ represents the strength of miRNA action. In the Dual degradation model the degradation constant of the mRNA-miRNA complex *k*_*rs*_ is introduced. Therefore, in short, the idea of an analysis given below is to fix a type of FFL and investigate how the target protein molecule number *p* will vary in all models considered.

#### Type 1 incoherent (1In) loop

This FFL is characterized by direct activation and indirect repression pathways, in which TF acts on target protein production.

*Variation of synthesis and degradation parameters*. Mathematical analysis showed that the increase of *h*_*p*_ in the Stop model and *h*_*g*_ coefficient in the Target degradation model results in the increase of the target protein quantity (Figure A2 of Additional file 2). The difference in target protein production is the largest for big values of these coefficients. The increase of the *k*_*rs*_ coefficient in the Dual degradation model results in the fall of the target protein quantity. Noteworthy, at early times (up to about *1000* seconds) the variation of *h*_*g*_ or *k*_*rs*_ coefficients has little influence on target protein production. In all models the difference in target protein production at different values of coefficients increases with time, as shown in Figure A2 of Additional file 2.

In all models the increase of *h*_*s*_ leads to increase in the target protein quantity (Figure 4). In both Target and Dual degradation models at early times (up to ca. *1000* seconds) the target protein production does not depend on *h*_*s*_ variation; at later times the difference between two bounding *h*_*s*_ values in the Target degradation model is much smaller than in other models. In all models the largest difference in target protein production is observed for large *h*_*s*_ values (Figure 4).

**Figure 4.**
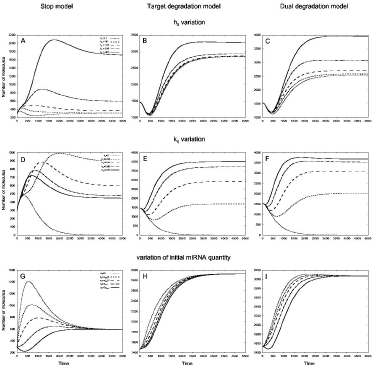
The behavior of 1*In* loop in response to variation of *h_s_, k_q_* and initial miRNA molecules numbers. The values of *h*_*s*_ coefficient defining the amount of TFs, at which the transcription rate of miRNA gene is half of its maximum value, were within 0 – 400 mol. interval, the translation rates *k*_*q*_ for TF were taken from 0 – 0.16*sec*^−1^ interval, initial quantities of miRNA were changed as described in section. Left column - Stop model, central column - Target degradation model, right column - Dual degradation model. **A - C**: In all models the quantity of target protein increases as *h*_*s*_ rise. **D**: In the Stop model the largest number of target protein molecules is observed at intermediate values of the *k*_*q*_ coefficient. **E -F**: In the Target and Dual degradation models *k*_*q*_ increase leads to increase of target protein quantity. **G and I**: In both Stop and Dual degradation models the form of target protein profile changes from the bell-shaped to the U-shaped one as the initial number of miRNA molecules rises. **H**: In the target degradation model all profiles show moderate dependence on the change of the initial number of miRNA molecules.

In all models the increase of *k*_*r*_ results in the increase of the target protein production, the effect of the *k*_*r*_ variation being larger at a later time (Figure A2 of Additional file 2). In all models the profiles nearly coincide up to a moment ca. 250 seconds and diverge afterwards. Later in the Stop model the profiles form a peak, which amplitude rises as *k*_*r*_ increases. Both the Target and Dual degradation models exhibit similar behavior: the profiles firstly tend to a minimum and increase to a constant value afterwards. In both models the steady state is achieved faster (around 2000 seconds) than in the Stop model (around 5000 seconds) (Figure A2 of Additional file 2).

In both the Target degradation and Dual degradation models applied to the 1*In* loop the increase of the TF translation rate *k*_*q*_ leads to increase of target protein quantity, the difference in target protein being the largest for small values of the coefficient (Figure 4). In Dual degradation model the steady state is reached earlier that in Target degradation model. In Stop model for this loop the largest number of target protein molecules is observed at *intermediate* values of the *k*_*q*_ coefficient (Figure 4).

*Variations in initial data*. In all models the variation of the initial number of TF and miRNA molecules does not change the number of target protein molecules at steady state. In both Stop and Dual degradation model the form of target protein profile changes from the *bell-shaped* to the *U-shaped* one as the initial number of miRNA molecules rises (Figure 4G, I). In Target degradation model all profiles have a form of increasing curve tending to a steady state and show moderate dependence on the change of the initial number of miRNA molecules (Figure 4H). The increase of the initial number of TF molecules results in the change of the target protein profile from the *U-shaped* form to the *bell-shaped* one in both the Target degradation and the Dual degradation models (Figure A2H, I of Additional file 2). In the Stop model all the target protein profiles have the *bell-shaped* forms, and their amplitudes decrease with the TF molecules number growth (Figure A2G of Additional file 2).

#### Type 1 coherent (1C) loop

This FFL is characterized by a synergetic action via both the direct and indirect pathways (see Figure 1).

*Variation of synthesis and degradation parameters*. The increase of both *h*_*p*_ and *h*_*g*_ coefficients results in the increase of the target protein quantity *p* (Figure A3 of Additional file 2). In the Stop model the difference in target protein production is the largest for large values of *h*_*p*_. In the Target degradation model only large values of *h*_*g*_ have a noticeable effect on the target protein production. The increase of *k*_*rs*_ in the Dual degradation model results in the fall down of the target protein quantity. At the very early times (up to about 500 seconds) the variations of either *h*_*g*_ or *k*_*rs*_ have a small influence on target protein production (Figure A3 of Additional file 2).

In all models for 1*C* loop the increase of *h*_*s*_ or *k*_*r*_ leads to increase in the target protein molecules number (Figure 5, Figure A3 of Additional file 2). In the first half of the whole time interval each profile forms a *peak*, which amplitude rises with *k*_*r*_ increase. In the Target and Dual degradation models the steady state quantities are reached faster than in the Stop model.

**Figure 5.**
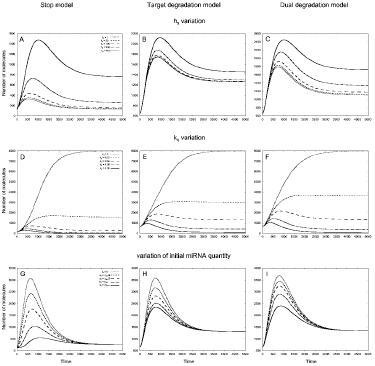
The behavior of 1*C* loop in response to variation of *h_s_, k_q_* and initial miRNA quantities. The parameter values are the same as of Figure 4. Left column - Stop model, central column - Target degradation model, right column - Dual degradation model. **A -C**: The dependence of target protein profiles on *h*_*s*_ variation. **D - F**: The dependence of target protein profiles on*k*_*q*_ variation. **G - I**: The dependence of target protein profiles on initial miRNA quantities. Initial quantities of miRNA were changed as described in section.

The increase of the TF translation rate *k*_*q*_ in all models applied to the 1*C* loop results in decrease of the target protein molecules number (Figure 5). Initially the difference in target protein production is not quite evident, later it becomes remarkable at small values of *k*_*q*_.

*Variations in initial data*. In all models for the 1*C* loop behavior the variations of the initial miRNA or TF molecule numbers do not influence the number of target protein molecules at steady state, see (Figure 5 and Figure A3 of Additional file 2). The increase in the initial number of miRNA molecules leads to the transformation of the *bell-shaped* target protein profile to the *U-shaped* profile in the Stop model (see Figure 5). In contrast to it, in the both Target degradation and Dual degradation models all profiles are bell-shaped, and their amplitudes decrease as the miRNA initial number increases (Figure 5). The gradual increase of the initial number of TF molecules results in the transformation of the bell-shaped target protein profile into uprising curve, tending to a constant value in all models (Figure A3 of Additional file 2).

#### Type 2 coherent (2C) loop

This FFL is characterized by coherent activation of a target via direct and indirect pathways (see Figure 1).

*Variation of synthesis and degradation parameters*. The increase of both *h*_*p*_ and *h*_*g*_ results in the increase of the target protein quantity *p* (Figure 6), while the increase of the miRNA-mRNA complex degradation coefficient *k*_*rs*_ leads to the opposite effect. In all models the profiles first tend to minima, then diverge. At very small time (up to about 500 seconds) the variation of the *h*_*g*_ or *k*_*rs*_ coefficients has a little influence on the target protein production. In all models the difference in target protein production is the largest for large values of the coefficients.

**Figure 6.**
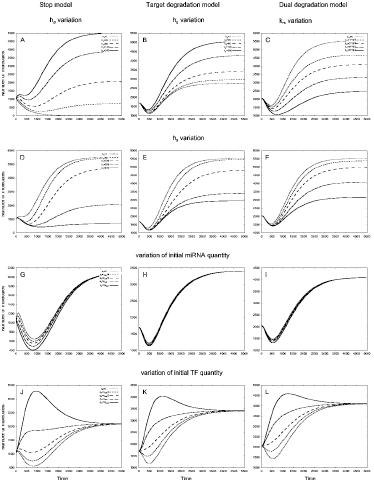
The behavior of 2*C* loop in response to variation of *h_p_, h_g_, k_rs_* and *h*_*s*_ coefficients, and initial miRNA and TF quantities. Parameters *h_p_, h_g_, k_rs_* define the action of miRNA on target mRNA, *h*_*s*_ is the dissociation coefficient. The miRNA and TF initial quantities were changed as described in section. Left column - Stop model, central column - Target degradation model, right column - Dual degradation model. **A**: In the Stop model the quantity of target protein increases as the value of *h*_*p*_ increases from 0 to 240 molecules. **B**: In the Target degradation model the quantity of target protein increases as the value of *h*_*g*_ increases from 0 to 240 molecules. C: The increase of the *k*_*rs*_ coefficient value from 0 to 8 *×* 10^−5^ *mol*^−1^ *sec*^−1^ results in the fall of the target protein quantity in the Dual degradation model. **D - F**: In all models the increase of *h*_*s*_ from 0 to 400 leads to decrease in target protein quantities. textbfG - I: The dependence of target protein quantities on initial miRNA molecules number in the models. **J -L**: The patterns of dependence of target protein quantities on initial TF quantities in the models.

In all models applied to the 2*C* loop the increase of dissociation constant *h*_*s*_ leads to decrease in target protein molecules number (Figure 6) and the largest difference in target protein quantities is observed at large values of this coefficient. In both Target and Dual degradation models the effect of *h*_*s*_ variation becomes especially evident at later times (after 1200 seconds).

In all models the increase of the maximal rate of transcription for target mRNA *k*_*r*_ results in the increase of the target protein quantity *p*, the effect of the *k*_*r*_ variation being larger at a later time (Figure A4 of Additional file 2). In the Stop model for this loop the target protein profiles firstly approach a maximum, then a minimum, and grow afterwards.

The increase of the TF translation rate *k*_*q*_ in all models of 2*C* loop leads to increase of target protein quantities, see Figure A4 of Additional file 2.

*Variation in initial data*. The variation of the initial miRNA and TF molecules number does not affect the steady state dynamics. In Stop model as the miRNA molecules initial number rises, the target protein profiles are transformed from wavelike to Ushaped pulse (Figure 6) and the difference in target protein dynamics corresponding to different initial miRNA quantities becomes almost negligible at later times. In both Target degradation and Dual degradation models the target protein profile has a form of U-shaped curve, however the difference between all the profiles is small at all times.

In all models of 2*C* loop the graph for the target protein quantity exhibits U-shaped profile when the initial number of TF molecules is small and bell-shaped profile when this number is large (Figure 6).

#### Type 2 incoherent (2In) loop

This FFL describes an indirect pathway of activation and direct pathway of the target repression (see Figure 1).

*Variation of synthesis and degradation parameters*. The increase of the *h*_*p*_ coefficient in the Stop model and *h*_*g*_ coefficient in the Target degradation model result in the target protein quantity growth. At very early time (up to about 500 seconds) the variation of *h*_*g*_ has small influence on target protein production, while at later time the larger values of *h*_*g*_ result in larger difference in target protein quantity (Figure A5 of Additional file 2). Growth of the miRNA-mRNA complex degradation coefficient *k*_*rs*_ results in fall of the target protein quantity in the Dual degradation model (Figure A5 of Additional file 2).

In all models for the 2*In* loop the increase of the *h*_*s*_ value results in decrease of the target protein molecules number (Figure A5 of Additional file 2). At small values of *h*_*s*_ the difference in target protein production is invisible in both Target and Dual degradation models at very early times and in all models at the second half of the whole time interval.

In all models the increase of the maximal rate for the target mRNA synthesis results in the increase of the target protein quantities, the effect of the *k*_*r*_ variation being larger at later time in the Stop model (Figure A5 of Additional file 2). For two other models the maximal effect of *k*_*r*_ variation is observed at the first half of the time interval. The pattern of dependence of the target protein profiles on *k*_*r*_ in the Stop model differs from that in two other models, namely, in the Stop model the target protein profile firstly approaches a maximum, than a minimum and grows afterwards (”wave-like” pattern), while in both the Target and the Dual degradation models the profiles form a peak (”pulse-like” pattern), which height is larger in the later model (Figure A5 of Additional file 2).

The increase of the rate *k*_*q*_ of the TF translation in the Target degradation and Dual degradation models of the 2*In* loop results in diminishment of the target protein molecules number (Figure A6 of Additional file 2). In the Stop model for this loop the largest number of target protein molecules at steady state is observed for *intermediate* values of *k*_*q*_, while the large values of *k*_*q*_ lead to almost cut off of protein production.

*Variations in initial data*. In all models the variation of initial number of TF or miRNA molecules does not change the number of target protein molecules at steady state. When the initial number of miRNA molecules gradually increases, the wave-like target protein profile is transformed to the U-shaped one in the Stop model applied to the 2*In* loop (Figure A6 of Additional file 2). In both Target degradation and Dual degradation models all profiles have bell-shaped form, and their amplitudes slightly decrease as the miRNA initial quantity grows.

In the Target and Dual degradation models for 2*In* loop the increase of initial number of TF molecules leads to the target protein profile changes from the bell-shaped to the U-shaped form (Figure A6 of Additional file 2). In the Stop model the variation of initial number of TF molecules weakly influences the target protein quantity: the profiles take a form of either wave-like or uprising curve and all of them tend to a steady state.

Our results show that FFLs mediated by miRNA may have many possible outcomes, depending on interaction between the loop elements. The target protein profiles can take different forms, which are unambiguously defined by initial conditions and model coefficients. In most cases the variation of model coefficients leads to results, which could be intuitively explained by consideration of the miRNA action in a model and the topology of a loop, however, in several cases the response of the system is hardly predictable. This especially concerns the variation of *k*_*q*_, the parameter, which defines the quantity of TF. In the Stop model in 1C loop the TF represses and in 2C loop it activates the target protein synthesis in parallel via direct or indirect pathways, respectively. Therefore in 2C loop the rise of *k*_*q*_ leads to an increase in target protein quantity, while in 1C loop the relation is opposite (Figure 5 and Figure A4 of Additional file 2). However, in the Stop model applied to both 1In and 2In loops the maximal number of the target protein molecules is observed at intermediate *k*_*q*_ values, moreover, in the last one the effect becomes visible only after 750 seconds (Figure 4 and Figure A6 of Additional File 2). In general, any noticeable variation in molecule quantity provided by an intermediate value of a parameter can be explained by simple mathematical analysis of the Hill function sigmoid, see Additional text 1.

### Noise buffering by miRNA

The comparative analysis performed above shows significantly diverse reaction of each FFL to variation of coefficients in each of the models considered.

In a wet lab it seems to be easier to identify a type of the FFL rather than to reveal the regulation details, which will be helpful in selection of the miRNA action model. The analysis of the temporal behavior of the FFL together with the ability to find out a unique solution for every set of parameters may provide the way to select the most probable mechanism of miRNA action if the type of FFL is known. However, a noise in data can corrupt an ideal behavior of FFL in absence of any perturbation.

It is widely believed that miRNA can buffer the consequences of noise in a cell. A simple mathematical model based on assumption that miRNA represses translation of its target was recently introduced to explore the ability of the 1*In* FFL to buffer fluctuations in upstream TF at steady state [30]. We demonstrated already that the behavior of a FFL is model-dependent, and it is reasonable to study the ability of FFLs to buffer fluctuations in upstream regulator quantity. on the assumption of the miRNA action, different from the translational repression. Besides, the results of our analysis also demonstrate an imperfection of approaches based on the analysis of model behavior at steady state only: as it is shown in Figures 5, 6 and A2, A3 of Additional file 2 the dynamical behavior of FFLs can be multivariant at early times even when the quantity of target protein at steady state is the same.

Consequently, we decide to investigate the ability of all FFLs with miRNA to buffer a noise caused by TF at all time moments and in all models. At each time moment we introduce the random fluctuations in TF quantity and measure how these fluctuations affect the target protein molecule number. We also consider how the variation of model coefficients influences the ability of the loops in noise damping. The efficiency of the FFLs in controlling the fluctuations of the target protein in response to noise introduced by TF depends on the number of TF molecules (which is a function of both *k*_*w*_ and *k*_*q*_), the number of miRNA copies (depending on *k*_*s*_ and *h*_*s*_) and the strength of miRNA action on target mRNA (defined by *h_p_, h_g_* and *k*_*rs*_ coefficients in the models). We studied the ability of FFLs to buffer noise as a function of each of these three quantities, changing a corresponding coefficient and keeping fixed all others. For each combination of a model, loop and coefficient value we performed *100* numerical simulation runs to estimate the average value of the parameter *ε* and the number of experiments with positive value of this parameter as described in section. Positive values of *ε* mean that a loop cannot dampen noise introduced by TF, and vice versa negative values of this coefficient testify the ability of the loop to reduce TF noise.

The 1 *In* FFL shows the best ability to buffer noise introduced by TF: the noise is strongly decreased at the level of target protein production in all models and within a wide range of parameter variation Figure 7, Table 1. In all models the noise buffering increases with *k*_*q*_ parameter for TF translation rate. At small values of *k*_*q*_ the Stop model is more effective in noise reduction than two other models, while at large values of this parameter all models show similar ability to reduce noise. The variation of parameters that define the miRNA level in a loop, namely *h*_*s*_ and *k*_*s*_, has small effect on ability of this loop to buffer noise in both the Target degradation and Dual degradation models. In the Stop model the maximal effect is achieved at *intermediate values* of these coefficients, the rate of noise reduction being the highest among all the models. Both in Target and Dual degradation models the ability of 1 *In* FFL to buffer noise does not significantly change when a parameter, which defines the strength of miRNA action *(h*_*g*_ or *k*_*rs*_), has large variation. In the Stop model the maximal effect is achieved at *intermediate values* of the *h*_*p*_ coefficient.

**Figure 7.**
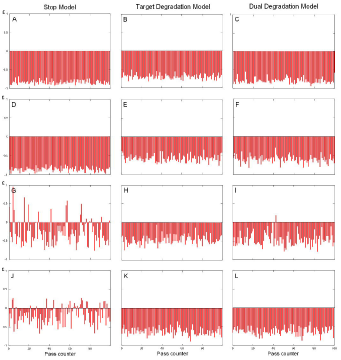
The ability of FFLs to buffer noise introduced by TF in frame of different models. In each panel the values of parameter *ε* calculated in 100 experiments are shown as narrow vertical lines. *ε* < 0 means that the noise is buffered in a loop, *ε* > 0 means non-ability of the loop to dampen noise. The coefficients and initial conditions for each experiment are given in section. **A -C**: Type 1 incoherent loop is able to buffer noise introduced by TF in all models. **D - F**: Type 2 incoherent loop buffers TF noise in all models. **G**: Type 1 coherent loop is a bad buffer in frame of the Stop model. **I, H**: Type 1 coherent loop is able to dampen TF noise in frame of the Target and Dual degradation models. **J**: Type 2 coherent loop is a bad buffer in frame of the Stop model. **K, L**: Type 2 coherent loop is able to buffer TF noise under the Target and Dual degradation models.

**Table 1.** The ability of 1*In* loop to buffer TF noise under variation of parameters in all three models.

| Parameter values | Stop Model |  | Degr. Model |  | Dual Degr. Model |  |
| --- | --- | --- | --- | --- | --- | --- |
| | $\bar{\varepsilon}$ | $n$ | $\bar{\varepsilon}$ | $n$ | $\bar{\varepsilon}$ | $n$ |
| $h_s = 1$ | -0.66 | 0 | $\{-0.67\}$ | 0 | -0.66 | 0 |
| $h_s = 100$ | **-0.84** | 0 | $\{-0.68\}$ | 0 | -0.69 | 0 |
| $h_s = 200$ | **-0.84** | 0 | -0.71 | 0 | $\{-0.74\}$ | 0 |
| $h_s = 400$ | -0.66 | 0 | -0.76 | 0 | $\{-0.75\}$ | 0 |
| $k_q = 0.02$ | -0.80 | 0 | -0.48 | 0 | -0.50 | 0 |
| $k_q = 0.04$ | -0.84 | 0 | -0.71 | 0 | -0.74 | 0 |
| $k_q = 0.08$ | -0.92 | 0 | -0.88 | 0 | -0.91 | 0 |
| $k_q = 0.16$ | -0.97 | 0 | -0.96 | 0 | -0.97 | 0 |
| $k_s = 0.01$ | -0.70 | 0 | $\{-0.71\}$ | 0 | -0.70 | 0 |
| $k_s = 0.25$ | **-0.89** | 0 | $\{-0.72\}$ | 0 | -0.72 | 0 |
| $k_s = 0.50$ | -0.84 | 0 | $\{-0.70\}$ | 0 | $\{-0.75\}$ | 0 |
| $k_s = 0.75$ | -0.80 | 0 | $\{-0.70\}$ | 0 | $\{-0.74\}$ | 0 |
| $h_p(h_g) = 30; k_{rs} = 1 \cdot 10^{-5}$ | -0.78 | 0 | -0.68 | 0 | $\{-0.73\}$ | 0 |
| $h_p(h_g) = 60; k_{rs} = 2 \cdot 10^{-5}$ | -0.83 | 0 | -0.70 | 0 | $\{-0.74\}$ | 0 |
| $h_p(h_g) = 120; k_{rs} = 4 \cdot 10^{-5}$ | **-0.90** | 0 | $\{-0.73\}$ | 0 | $\{-0.76\}$ | 0 |
| $h_p(h_g) = 240; k_{rs} = 8 \cdot 10^{-5}$ | -0.82 | 0 | $\{-0.73\}$ | 0 | $\{-0.77\}$ | 0 |

For each model and for each parameter value we present the average $\varepsilon$ value (left column) and the number $n$ of experiments with *positive* value of this coefficient (right column). The method for calculation of $\varepsilon$ is described in section . Negative $\varepsilon$ values mean that TF noise is dampen in a loop. The extremal $\varepsilon$ values achieved at intermediate parameter values are shown in bold. The Wilcoxon Rank-Sum Test was used to test for significance of difference between the values of $\varepsilon$ for adjacent parameter values. The differences which are statistically insignificant at the $\alpha = 0.05$ level are placed in parentheses.

The ability of the *2In* loop to decrease noise introduced by TF essentially depends on the *k*_*q*_ coefficient values: all the models are not able to efficiently buffer noise for large values of this coefficient (at *k*_*q*_ *=* 0.16 the *ε* coefficient was positive in 9 – 14% of experiments), however, in the Stop and Target degradation models this effect becomes evident at larger *k*_*q*_ values that in the other model (Table 2). In the Target degradation and Dual degradation models the variation of other coefficients has small effect on the ability of the loop to reduce noise. It is worth to note, that for larger values of both *k*_*s*_ and *h*_*s*_ the Stop model is more effective in noise buffering, than two other models. In frame of the Stop model the higher are *h*_*s*_ values, the higher is the ability of *2In* loop to reduce TF noise. On the contrary, the strongest noise reduction in this loop is achieved at intermediate values of the *k*_*s*_ and *h*_*p*_ coefficients.

**Table 2.** The ability of 2*In* loop to buffer TF noise under variation of parameters in all three models.

| Parameter values | Stop Model |  | Degr. Model |  | Dual Degr. Model |  |
| --- | --- | --- | --- | --- | --- | --- |
| | $\bar{\varepsilon}$ | $n$ | $\bar{\varepsilon}$ | $n$ | $\bar{\varepsilon}$ | $n$ |
| $h_s = 1$ | -0.50 | 0 | -0.49 | 0 | -0.48 | 0 |
| $h_s = 100$ | -0.65 | 0 | **-0.57** | 0 | -0.54 | 0 |
| $h_s = 200$ | -0.81 | 0 | -0.54 | 0 | **-0.56** | 0 |
| $h_s = 400$ | -0.84 | 0 | -0.48 | 0 | -0.52 | 0 |
| $k_q = 0.02$ | $\{-0.82\}$ | 0 | -0.71 | 0 | -0.76 | 0 |
| $k_q = 0.04$ | $\{-0.82\}$ | 0 | -0.53 | 0 | -0.56 | 0 |
| $k_q = 0.08$ | -0.56 | 0 | -0.44 | 0 | $\{-0.39\}$ | 5 |
| $k_q = 0.16$ | -0.35 | 9 | -0.33 | 13 | $\{-0.33\}$ | 14 |
| $k_s = 0.01$ | -0.47 | 0 | -0.48 | 0 | $\{-0.49\}$ | 0 |
| $k_s = 0.25$ | -0.73 | 0 | **-0.57** | 0 | $\{-0.51\}$ | 0 |
| $k_s = 0.50$ | **-0.82** | 0 | $\{-0.54\}$ | 0 | $\{-0.56\}$ | 0 |
| $k_s = 0.75$ | -0.71 | 0 | $\{-0.51\}$ | 0 | $\{-0.57\}$ | 0 |
| $h_p(h_g) = 30; k_{rs} = 1 \cdot 10^{-5}$ | -0.65 | 0 | -0.48 | 0 | $\{-0.53\}$ | 0 |
| $h_p(h_g) = 60; k_{rs} = 2 \cdot 10^{-5}$ | **-0.81** | 0 | $\{-0.54\}$ | 0 | $\{-0.55\}$ | 0 |
| $h_p(h_g) = 120; k_{rs} = 4 \cdot 10^{-5}$ | -0.74 | 0 | $\{-0.57\}$ | 0 | -0.58 | 0 |
| $h_p(h_g) = 240; k_{rs} = 8 \cdot 10^{-5}$ | -0.57 | 0 | -0.53 | 0 | -0.63 | 0 |

For each model and for each parameter value we present the average $\varepsilon$ value (left column) and the number $n$ of experiments with *positive* values of this coefficient (right column). The method for calculation of $\varepsilon$ is described in section . Negative $\varepsilon$ mean that TF noise is dampen in a loop. The extremal $\varepsilon$ values achieved at intermediate parameter values are shown in bold. The Wilcoxon Rank-Sum Test was used to test for significance of difference between the $\varepsilon$ values for adjacent parameter values. The differences which are statistically insignificant at the $\alpha = 0.05$ level are placed in parentheses.

Both *1C* and *2C* loops are bad buffers in frame of the Stop model but generally reduce the TF noise under both the Target and Dual degradation models (Figure 7). In the frame of the Target and Dual degradation models the variation of all coefficients (except of *k*_*q*_ in both loops) exposes to the ability of these loops to buffer noise weaker than in the Stop model (Tables 3 and 4). In this model the behavior of the *2C* loop and, to smaller extent, of the *1C* loop shows a strong dependence on parameters. Below we address the ability of coherent loops to buffer noise in detail.

**Table 3.**
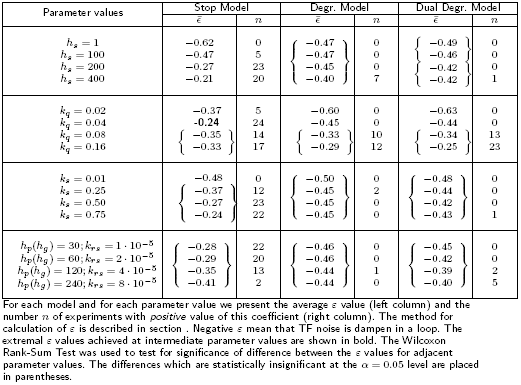
The ability of 1*C* loop to buffer TF noise under variation of parameters in all three models.

In the Stop model the *1C* loop is inefficient in reduction of TF noise at any value of the *k*_*q*_ coefficient, showing the worst reduction at intermediate values (*k*_*q*_=0.04) (Table 3). For large *k*_*q*_ values (*k*_*q*_=0.08 and higher) and in both the Target and Dual degradation models this loop loses the ability to efficiently reduce noise and starts to deal with the noise as in the Stop model.

The *1C* loop is able to effectively buffer noise only for very small values of *h*_*s*_ and *k*_*s*_ coefficients in the Stop model: the larger are these coefficients, the less efficient is the noise reduction (Table 3). The loop shows non-ability to reduce noise in small number of experiments at *h*_*s*_ *=* 400 in both Target and Dual degradation models, as well as at intermediate *k*_*s*_ *=* 0.25 value in the Target degradation model and at *k*_*s*_ *=* 0.75 in the Dual degradation model.

In the Stop model the *h*_*p*_ increase improves the ability of the loop to buffer noise as both the average *ε* value and the number of experiments, in which TF noise was not dampen (positive *ε* value) decrease (Table 3). In the Dual degradation model the *1C* loop shows non-ability to buffer TF noise in 2-5% of experiments.

The ability of *2C* loop to reduce TF noise increases with the value of *k*_*q*_ in the Stop model (Table 4). For large values of it (*k*_*q*_=0.08 and higher) the TF noise was reduced in all numerical simulations. In two other models the TF noise is always dampen (with one exception of *k*_*q*_ *=* 0.02 in the Dual dagradation model, see (Table 4), however the efficiency of noise buffering increases as the value of the *k*_*q*_ rises.

**Table 4.**
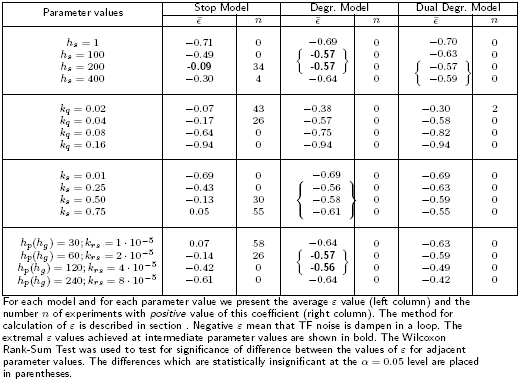
The ability of 2*C* loop to buffer TF noise under variation of parameters in all three models.

The dependence of the efficiency of noise damping on *h*_*s*_ variation in the *2C* loop and in frame of Stop model is complex: noise is efficiently reduced for very small *h*_*s*_ values, while the efficiency of noise reduction decreases as the *h*_*s*_ value grows, however the worst noise reduction happens at intermediate *h*_*s*_ values (*h*_*s*_=200). In the Stop model the ability of *2C* loop to buffer noise decreases with the growth of *k*_*s*_: at small *k*_*s*_ *≤* 0.25 values the noise damping was observed in all simulations, while for higher *k*_*s*_ values the fluctuations of the target protein molecules number are not reduced in many simulation runs (Table 4).

The *2C* loop is unable to effectively buffer TF noise in all experiments when the values of the *h*_*p*_ coefficient are small (below or equal to 60), at larger coefficient values this loop starts to efficiently buffer noise (Table 4). The noise is effectively reduced in the Dual degradation model of *2C* loop for all values of the *k*_*rs*_ coefficient, however the ability of dampening decreases as the value of this coefficient rises.

## Conclusions

We performed here the theoretical analysis of a gene network sub-system, containing a FFL mediated by TF and miRNA. We have shown that different mechanisms of miRNA action lead to a variety of types of dynamical behavior of FFLs during cell cycle and govern their ability to dampen noise caused by TF fluctuations.

The molecular mechanisms of miRNA action are not clear so far, and we elaborate three mathematical models introduced in [30], that describe the gene expression in miRNA mediated FFL under the assumption of different mechanisms of miRNA action. In the Stop model miRNA represses translation of its target mRNA, in the Target degradation model miRNA promotes the target degradation, and in the Dual degradation model miRNA is not re-used, but degrades along with target mRNA.

Due to an intrinsic complexity and non-linearity of biological systems it is a hard task to obtain any analytic solution to differential equations, which describe the regulation in FFL with miRNA under the models considered. These equations are non-linear, fortunately we were able to obtain the exact solutions to some of them, namely, to those describing target mRNA and miRNA production in the Stop model and for several biologically relevant values, i.e.,

- for slow degradation of miRNA (when miRNA is degraded two times slower than TF or its mRNA),
- fast degradation of miRNA (when miRNA is degraded two times faster than TF or its mRNA) and
- very fast degradation of miRNA (when miRNA is degraded three times faster than TF) (see Additional file 1 for details).

Despite of the fact that miRNAs are generally stable molecules, it was shown recently that individual miRNAs may be exposed to an accelerated decay [18].

We used the exact solutions to check the results of numerical simulations obtained for all other coefficient sets and initial conditions, and proven the validity of these results. In general, the exact solutions obtained could be used as a genuine check point in numerical simulations of larger networks with miRNA embedded into a FFL motif.

We have rigorously proven the uniqueness of solutions to all the models under consideration, i.e., in the models considered there is the one-to-one correspondence between the given parameter set and the solution, describing the dynamics of target protein production in FFL.

Study of cell components behaviour at steady state is conventional, however it is worth to note that the steady states are not completely informative even in non-biotic systems, consisting of almost identical clusters of several atoms/molecules. In biology the steady states are not unique: very different pathways may lead to the same stationary position. From the general viewpoint of dynamic control theory an early time seems to be the most promising one for tentative control/influence onto the loop dynamics, either by noise or by any external factor. That is why we first examined the FFL dynamics quantitatively, and at the whole time interval of cell cycle, alternatively to recent qualitative consideration [30].

Our results show that FFLs mediated by miRNA and TF may have many possible outcomes, depending on interaction between the loop elements. The target protein profiles can take different forms, which are unambiguously defined by initial conditions and model coefficients. This can be illustrated, when we consider the behavior of one and the same FFL under different models, and when both initial conditions and the level of a target protein at steady state are the same. Due to the difference in mechanisms of miRNA action the behavior of a FFL in time will be quite different in frame of different models. This situation is reconstructed for the 1*C* loop in Figure 8. It is evident that the behavior of the models at early times is different: the maximum of target protein production is the smallest in the Dual degradation model, while the time, at which this protein reaches the steady state is the largest one for the Stop model. It is noteworthy that these graphs correspond to models with different degradation coefficients of the target mRNA and protein, i.e. to different biological situations.

**Figure 8.**
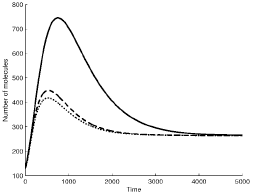
The target protein profiles in 1*C* FFL in different models. Both initial conditions and the steady state level of a target protein are identical. Solid line - Stop model, dashed line -Target degradation model, dotted line - Dual degradation model.

Contemporary experimental set up seems not be able to capture many facets of miRNA function. Indeed, in spite of a bunch of publications, describing the crucial role of miRNA in control of many biological processes and in progression of various diseases, the molecular mechanisms of miRNA action are still not evident [14, 15]. In a wet lab it is easier to identify a type of FFL, rather than to reveal the details of regulation, which will be helpful in selection of the miRNA action model. The analysis of the temporal behavior o f the FFL together with the ability to find out a unique solution for any set of parameters might help to select the most feasible mechanism of miRNA action for the type of FFL given.

The results obtained allow us to propose the following strategy for an ideal FFL: let us consider how a miRNA acts, having an information about the FFL topology and a given set of quantitative measurements of each player in the loop. If the initial quantities of molecules for each player in FFL are not known, we should firstly determine them in experiments. Using our analytical results, we shall be able to calculate in detail the temporal dependence for each player in the FFL and in each possible model for various values of coefficients. Next, we identify that graph, which passes through the given set of the (molecule quantities) points among all others. By virtue of the uniqueness theorem this graph will correspond to most feasible regulation type in the FFL among all models considered.

However FFLs are subjected to noise influence, and for this reason we studied the noise buffering in both coherent and incoherent FFLs, that led to conclusion that incoherent FFLs are better noise buffers than coherent ones (Figure 7). Moreover, even the parameter variation does not seriously affect the noise buffering ability of incoherent FFLs. Therefore the selection strategy proposed seems to be suitable to predict a model for incoherent FFLs.

The FFL dynamic behavior analysis performed for different models and parameter sets shows that an extremal value of target protein quantity (max or min, depending on a loop) may be accomplished for intermediate values of coefficients. This is valid also for the noise buffering problem, that required a simple analysis of the Hill sigmoid behavior, see Additional file 1.

In general, an action of any disturbance, e.g., a noise, having sufficiently small amplitude, (ca. 10 – 15% ; otherwise it would prevail over a signal in FFL) depends on the Hill function type and can be described as follows. For the Hill sigmoid functions, governing either an activation or a repression, the noise will be recognizable for intermediate molecule quantity. By virtue of analysis in the whole time interval we found that when the noise amplitude is high, but the total amount (the regular one plus the noise associated fluctuations) of miRNA is intermediate (the Hill function is far from any of almost constant limits), then the noise is translated directly into the protein production process and will *not be suppressed* by a loop.

When the noise amplitude is still high, however, the total amount of miRNA is sufficiently high, too (the Hill function is close to any of limits), then the noise level is relatively small to disturb the protein production and will *be suppressed* by a loop.

## Methods

Many details of mathematical analysis are given in Section and in the Supporting Information files; here we briefly describe the numerical simulations. The coefficients used in mathematical modeling were mostly taken from [30], and for the Stop model they are, as follows:

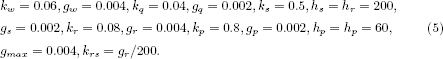

The other coefficients used in both the Degradation and the Dual degradation models are:

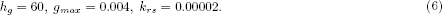

Numerical simulation of dynamics in each loop was made for the time period of *t = 5000* seconds. The initial numbers of components in each loop were equal to one half of their steady state values.The dynamics of solutions at early stages depends not only on coefficients, but on initial values, too. To analyze how the quantity of target protein molecules depends on the numbers of TF and miRNA molecules at initial time moment these numbers were changed from 0 to one quarter and one half, as well as to the values of two and four times higher, than at steady state (in normalized quantities).

For the noise simulations we introduced a vector *z* with the correlation function *K*(*τ*) = exp(–|*τ*|), that was obtained in a form:

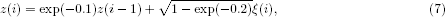

where *ξ* is another vector, obtained by the MATLAB procedure, which contains the Gaussian white noise. The vector *z* represents the model of a stationary Markov process, and we may add its values at the moments *t*_*i*_ to the number of molecules of TF, which is used in equations for the quantities of miRNA and target mRNA, and solve these equations numerically. In fact, the procedure mentioned is based on the Uhlenbeck-Ornstein process (1930), the only one, which is stationary random, Gaussian and Marcovian simultaneously. The process is widely used now in mathematical modelling of noise instead of any numerical version of the so called “ white noise”. The particle velocity in this process is finite, and taking into account the difference between a random force and the idealised white noise, one may provide a finite acceleration of the particle, too.

We calculated the numbers *q* of TF molecules, of miRNA (*s*), of the target mRNA (*r*) and of target protein (*p*) with the noise component and without it, and afterwards we found the values *N_A_ = max_t_*|*q(t) – q_n_(t)|, N_B_ = max_t_*|*p(t) – p*_*n*_*(t)*| to estimate the relative power of noise. Here *q(t), p(t)* are the numbers of molecules of transcription factor and target protein without the noise component, respectively, while *q_n_(t), p_n_(t)* are corresponding quantities with noise. Therefore we may find the value of the estimation parameter *ε* :

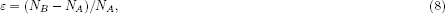

to conclude whether the relative noise level in target protein quantity is higher or lower, than in TF. Consequently, *ε <* 0 means that the noise level in target protein is lower than in TF, and the noise is buffered in the loop.

We performed calculations with 100 different noise vectors *z* for each loop and each model, estimated *ε* in every calculation, studied and calculated an average value of *ε*, as well as the number of calculations with positive *ε* value. We used the Wilcoxon test for these averaged values of *ε* to compare results for the same loops and same models under variation of the model coefficients.

## Author Contributions

AS and MS proposed and designed the study, MD and AS obtained all the mathematical solutions to the problem, MD and MS analyzed the modeling results. AS and MS wrote and typeset the manuscript with considerable input from MD.

## Acknowledgements

We are thankful to Sergey Berezin for valuable advises.This work is partly supported by RFBR grants 14-01-00334, 14-04-01522, EC collaborative project Health-F5-2010-260429 SysPatho and the Russian Federation Ministry of Education and Science program 5-100-2020.

## Additional information

Additional file 1 (pdf) contains details of mathematical analysis of the coupled ODE

Additional file 2 (pdf) contains several figures useful for better understanding the results presented in the main text.

## References

1. He, L., Hannon, G.J.: MicroRNAs: small RNAs with a big role in gene regulation. Nat Rev Genet 5, 522–531 (2004)

2. D, H.B.: MicroRNAs in vertebrate development. Curr Opin Genet Dev. 15(4), 410–415 (2005)

3. Bushati, N., Cohen, S.M.: microRNA functions. Annu Rev Cell Dev Biol 23, 175–205 (2007)

4. Avraham, R., Yarden, Y.: Regulation of signalling by microRNAs. Biochem. Soc. Trans. 40(1), 26–30 (2012)

5. Calin, G.A., Ferracin, M., Cimmino, A., Di Leva, G., Shimizu, M., Wojcik, S.E., Iorio, M.V., Visone, R., Sever, N.I., Fabbri, M., Luliano, R., Palumbo, T., Pichiorri, F., Roldo, C., Garzon, R., Sevignani, C., Rassenti, L., Alder, H., Volinia, S., Liu, C.G., Kipps, T.J., Negrini, M., Croce, C.M.: A microRNA signature associated with prognosis and progression in chronic lymphocytic leukemia. N Engl J Med 353, 1793–1801 (2005)

6. Alvarez-Garcia, I., Miska, E.A.: MicroRNA functions in animal development and human disease. Development 132, 4653–4662 (2005)

7. Beezhold, K.J., Castranova, V., Chen, F.: Microprocessor of microRNAs: regulation and potential for therapeutic intervention. Mol Cancer 9, 134 (2010)

8. Flynt, A.S., Lai, E.C.: Biological principles of microRNA-mediated regulation: shared themes amid diversity. Nature Reviews Genetics 9(11), 831–42 (2008)

9. Enright, A.J., John, B., Gaul, U., Tuschl, T., Sander, C., Marks, D.S.: MicroRNA targets in drosophila. Genome Biol 5(1), 1 (2003)

10. Brennecke, J., Stark, A., Russell, R.B., Cohen, S.M.: Principles of microRNA-target recognition. PLoS Biol 1(1), 13 (2005)

11. Grun, D., Wang, Y.-L., Langenberger, D., Gunsalu, K.C., Rajewsky, N.: microRNA target predictions across seven drosophila species and comparison to mammalian targets. PLoS Comput. Biol. 1(1), 13 (2005)

12. Hornstein, E., Shomron, N.: Canalization of development by microRNAs. Nat Genet 38, 20–24 (2006)

13. Friedman, R.C., Farh, K.K.-H., Burge, C.B., Bartel, D.P.: Most mammalian mRNAs are conserved targets of microRNAs. Genome Research 19(1), 92–105 (2008)

14. Baek, D., Vill´en, J., Shin, C., Camargo, F.D., Gygi, S.P., Bartel, D.P.: The impact of microRNAs on protein output. Nature 455(7209), 64–71 (2008)

15. Selbach, M., Schwanh¨ausser, B., Thierfelder, N., Fang, Z., Khanin, R., Rajewsky, N.: Widespread changes in protein synthesis induced by microRNAs. Nature 455(7209), 58–63 (2008)

16. Bazzini, A.A., Lee, M.T., Giraldez, A.J.: Ribosome profiling shows that miR-430 reduces translation before causing mRNA decay in zebrafish. Science 336(6078), 233–7 (2012)

17. Guo, H., Ingolia, N.T., Weissman, J.S., Bartel, D.P.: Mammalian microRNAs predominantly act to decrease target mRNA levels. Nature 466, 835–841 (2010)

18. Ruegger, S., Grosshans, H.: Micro rna turnover: when, how, and why. Trends Biochem Sci. 37(10), 436–446 (2012)

19. Baccarini, A., Chauhan, H., Gardner, T.J., Jayaprakash, A.D., Sachidanandam, R., Brown, B.D.: Kinetic analysis reveals the fate of a microRNA following target regulation in mammalian cells. Current Biology 21(5), 369–376 (2011)

20. Shen-Orr, S.S., Milo, R., Mangan, S., Alon, U.: Network motifs in the transcriptional regulation network of escherichia coli. Sciencet 31(1), 64–68 (2002)

21. Milo, R., Shen-Orr, S.S., Itzkovitz, S., Kashtan, N., Chklovskii, D., Alon, U.: Network motifs: simple building blocks of complex networks. Nat Genet 298, 824–827 (2002)

22. Alon, U.: An introduction to systems biology. Design principles of biological circuits, p. 301. Chapman& Hall / CRC, ??? (2006)

23. Yu, X., Lin, J., Zack, D.J., Mendell, J.T., Qian, J.: Analysis of regulatory network topology reveals functionally distinct classes of microRNAs. Nucleic Acids Research 36, 6494–6503 (2008)

24. Herranz, H., Cohen, S.M.: MicroRNAs and gene regulatory networks: managing the impact of noise in biological systems. Genes & Development 24(13), 1339–44 (2010)

25. Stark, A., Brennecke, J., Bushati, N., Russell, R.B., Cohen, S.M.: Animal MicroRNAs confer robustness to gene expression and have a significant impact on 3’UTR evolution. Cell 123(6), 1133–1146 (2005)

26. Farh, K.K., Grimson, A., Jan, C., Lewis, B.P., Johnston, W.K., Lim, L.P., Burge, C.B., Bartel, D.P.: The widespread impact of mammalian microRNAs on mRNA repression and evolution. Science 310, 1817–1821 (2005)

27. Sood, P., Krek, A., Zavolan, M., Macino, G., Rajewsky, N.: Cell-type-specific signatures of microRNAs on target mRNA expression. Proc. Natl. Acad. Sci. USA 103, 2746–2751 (2006)

28. Giraldez, A.J., Mishima, Y., Rihel, J., Grocock, R.J., Van Dongen, S., Inoue, K., Enright, A.J., Schier, A.F.: Zebrafish mir-430 promotes deadenylation and clearance of maternal mRNAs. Science 312, 75–79 (2006)

29. Li, X., Cassidy, J.J., Reinke, C.A., Fischboeck, S., Carthew, R.W.: A microRNA imparts robustness against environmental fluctuation during development. Cell 137(2), 273–282 (2010)

30. Osella, M., Bosia, C., Corá, D., Caselle, M.: The role of incoherent microRNA-mediated feedforward loops in noise buffering. PLoS Computational Biology 7(3), 1001101 (2011)

31. Schwanhausser, B., Busse, D., Li, N., Dittmar, G., Schuchhardt, J., Wolf, J., Chen, W., Selbach, M.: Global quantification of mammalian gene expression control. Nature 473(7347), 337–342 (2011)

32. Li, J.J., Bickel, P.J., Biggin, M.D.: System wide analyses have underestimated protein abundances and the importance of transcription in mammals. PeerJ 2, 270 (2014)

33. von Dassow, G., Meir, E., Munro, E.M., Odell, G.M.: The segment polarity network is a robust developmental module. Nature 406, 188–192 (2000)

34. Kitano, H.: Biological robustness. Nat Rev Genet 5(11), 826–837 (2004)

